# Prior parental experience attenuates hormonal stress responses and alters hippocampal glucocorticoid receptors in both sexes of the biparental rock dove

**DOI:** 10.1101/2022.07.25.501460

**Authors:** Victoria S. Farrar, Jaime Morales Gallardo, Rebecca M. Calisi

## Abstract

In the face of challenges, animals must balance investments in reproductive effort versus their own survival. Physiologically, this tradeoff may be mediated by glucocorticoid release by the hypothalamic-pituitary-adrenal (HPA) axis and prolactin release from the pituitary to maintain parental care. The degree to which animals react to, and recover from, stressors likely affects their ability to maintain parental behavior and ultimately, fitness. However, less is known about how the stress response changes when animals gain parental experience, and what mechanisms may underlie any effect of experience on hormonal stress responses. To address these questions, we measured the corticosterone (CORT) and prolactin (PRL) stress response in both sexes of the biparental rock dove (*Columba livia*) that had never raised chicks versus birds that had fledged at least one chick. We measured both CORT and PRL at baseline and after an acute stressor (30 minutes restraint). We also measured negative feedback ability by administering dexamethasone, a synthetic glucocorticoid that suppresses CORT release, and measuring CORT and PRL after 60 minutes. All hormones we measured when birds were not actively nesting, allowing us to assess any lasting effects of parental experience beyond the parental care period. Birds with parental experience had lower stress-induced and negative-feedback CORT, and higher stress-induced PRL than inexperienced birds. In a separate experiment, we measured glucocorticoid receptor subtype expression in the hippocampus, a key site of negative feedback regulation. We found that experienced birds expressed higher glucocorticoid receptors than inexperienced controls, which may mediate their ability to attenuate the hormonal stress response. Together, these results shed light on potential mechanisms by which gaining experience may improve parental performance and fitness.

**Summary statement:** Experienced rock dove parents show lower corticosterone and higher prolactin levels after an acute stressor than birds without parental experience and increased glucocorticoid receptor expression in the hippocampus may mediate this effect.

## Introduction

Following life-history theory, breeding animals can maximize fitness by prioritizing resource allocation towards reproductive efforts, such as parental care of their current brood, at a cost to personal survival, self-maintenance, and growth (Stearns, 1976; Williams, 1966). However, when faced with predation, food limitation, inclement weather, or social challenges, animals may enter an emergency life-history stage (Wingfield et al., 1998) and abandon the current reproductive effort in order to survive (Wingfield and Sapolsky, 2003). Much research in recent decades has been on the physiological mechanisms underlying these tradeoffs (Ricklefs and Wikelski, 2002; Zera and Harshman, 2001), especially in the face of stressors (Romero and Wingfield, 2016).

Endocrine mechanisms, specifically glucocorticoid hormones (corticosterone or cortisol; CORT) and prolactin (PRL), have been strongly implicated in tradeoffs between survival and reproduction due to their pleiotropic effects on energetic state, metabolism, and reproduction. In response to challenges, the hypothalamus releases corticotropin-releasing factor (CRF), which stimulates the pituitary to release adrenocorticotropic hormone (ACTH), which leads to CORT synthesis and release from the adrenal glands (Aguilera, 2016). This hypothalamic-pituitary-adrenal (HPA) endocrine axis is relatively conserved across vertebrates (Blas, 2015; Romero and Gormally, 2019). Increased CORT can promote survival during challenges by increasing glucose availability via gluconeogenesis, mobilizing free fatty acids as an energy source and potentiating foraging and escape behaviors (Landys et al., 2006; Sapolsky et al., 2000; Wingfield et al., 1998); but see (Taff et al., 2022). Baseline CORT can also increase naturally during energetically-costly stages, like breeding (Bonier et al., 2011; Romero, 2002). However, elevated CORT in the face of stressors can also directly inhibit reproductive physiology and behavior, including parental behavior (Wingfield and Sapolsky, 2003). Conversely, elevated PRL from the pituitary promotes resource allocation towards parental efforts in vertebrates, by facilitating lactation, offspring attendance and provisioning (as examples; (Buntin, 1996; Freeman et al., 2000; Smiley, 2019). In a stress context, reduced PRL may reduce investment away from parental effort and behavior in birds (the “prolactin stress hypothesis”, (Angelier and Chastel, 2009; Chastel et al., 2005). However, acute stress often leads to increased PRL in mammals (Torner, 2016), so the prolactin stress hypothesis may not generalize across vertebrates. Nonetheless, the CORT and PRL stress responses can yield important insights into the tradeoff between survival and energetic balance versus reproductive effort (Angelier et al., 2016; Angelier and Chastel, 2009) when measured together within individuals.

However, less is known about how previous parental experience may affect these hormonal stress responses. Young, inexperienced individuals may have constrained physiological abilities to modulate hormones in response to stress (the “constraint hypothesis”; Curio, 1983), or they may restrain from modulating such responses due to relatively higher future reproductive opportunities (“restraint hypothesis”)(Curio, 1983). Studies in long-lived seabirds suggest that age may lead to attenuated CORT and PRL stress responses, where older individuals show lower stress-induced CORT and higher stress-induced PRL (Heidinger et al., 2010, 2006). In contrast, other studies found no effects of age on stress-induced CORT, but did find older seabirds maintained higher PRL levels at baseline or after stress (Angelier et al., 2007a, 2007b). As gaining breeding experience necessarily requires time that ages individuals, any effects of age on stress responses seen in these studies may be modulated in part by parental experience. Indeed, previous breeding experience in these long-lived seabirds may be a better predictor of baseline CORT and PRL levels than age alone (Angelier et al., 2007b, 2006). Baseline PRL levels have also been shown to increase with subsequent breeding experiences within individuals (Smiley and Adkins-Regan, 2016). However, whether parental experience itself alters hormonal stress responses when animals are relatively similar in age remains unclear.

Upstream of hormone release, neural receptors densities may also underlie differences in hormonal stress responses that may appear with breeding experience. Prior breeding experience has been shown to affect endocrine systems, such as pituitary prolactin cell counts or neural prolactin receptors (Anderson et al., 2006; Christensen and Vleck, 2015; Farrar et al., 2022), but effects on glucocorticoid-specific regulation remain unstudied. CORT exerts effects through two genomic receptor types, the high-affinity mineralocorticoid receptors (Type I; *MR*) the lower-affinity glucocorticoid receptors (Type II; *GR*), as well as membrane-based receptors (Breuner and Orchinik, 2009). These genomic receptors are hypothesized to play distinct roles, where the high-affinity MR enacts permissive effects of CORT at baseline levels, and the lower-affinity GR enacts suppressive and adaptive actions in response to elevated CORT levels, such as those seen after stressors (Romero, 2004; Sapolsky et al., 2000). While these receptors are found throughout the body, hippocampal *MR* and *GR* may be especially important for negative feedback of CORT after a stressor. Both hippocampal *MR* and *GR* have been shown to mediate hypothalamus-pituitary-adrenal axis activity and CORT release in mammals (de Kloet et al., 1998; Jacobson and Sapolsky, 1991; R. de Kloet and C. Meijer, 2019), though evidence is limited in birds (Smulders, 2017). The balance of these receptor subtypes may also play a role in maintaining homeostasis and avoiding stress pathology. For example, reduced hippocampal *GR* expression led to increased CORT levels after restraint stress in transgenic mice, presumably due to reduced negative feedback inhibition, but overexpressed *MR* with reduced *GR* undid this effect (Harris et al., 2013). In birds, hippocampal glucocorticoid receptor expression can change during seasonal or breeding transitions (Krause et al., 2015; Lattin and Romero, 2013). However, no study to our knowledge has evaluated how prior parental experience may alter hippocampal glucocorticoid receptor expression while also measuring animals’ ability to negatively feedback CORT levels.

To address these questions, we first examined hormonal stress responses in CORT and PRL in non-actively-nesting rock doves (*Columba livia*) that differed in prior parental experience with chicks. We used dexamethasone, a synthetic glucocorticoid, to induce maximal negative feedback when collecting stress series (an established method in avian endocrinology; (Lattin and Kelly, 2020), allowing us to compare baseline, stress-induced, and negative feedback levels of each hormone. In a second experiment, we extended our analysis into the brain, where we measured hippocampal gene expression of *MR* and *GR* using quantitative PCR. By capitalizing on a captive breeding population of biparental rock doves, we were able to collect data from individuals of both sexes with known breeding histories and ages.

We aimed to test a variation of the “constraint” and “restraint” hypotheses (Curio, 1983), modified for the effects of parental experience (instead of age alone). We hypothesized that prior parental experience would lead to reduced constraint on hormonal modulation, thus improving the ability to attenuate stress responses and invest in reproduction. Accordingly, we predicted that birds with prior experience with chicks would have lower CORT and higher PRL after an acute stressor and after negative feedback than birds that had never previously raised chicks. Further, we hypothesized that parental experience alters hippocampal glucocorticoid receptors, enabling more flexible hormonal stress responses. We then predicted that birds who had raised chicks would have higher hippocampal *GR* and/or *MR* expression than inexperienced birds. As both sexes of rock doves participate nearly equally in incubation and chick care (Abs, 1983; Johnston, 1992), we predicted we would observe no sex differences in hormone stress responses or glucocorticoid receptor subtypes.

## Methods

### Experiment 1: Hormonal responses to stress

#### Subjects and study design

We collected stress series (consisting of three blood samples) from 35 adult rock doves (*Columba livia*) of both sexes between March and June 2021. All subjects were born in captivity and housed in a semi-natural, social aviary environment. Each outdoor flight aviary (1.5 × 1.2 × 2.1m) is exposed to ambient temperatures and natural daylight, which is supplemented with artificial lighting on a 14L:10D photoperiod. Birds are provided *ad libitum* food (whole corn and turkey/game bird protein starter, 30% protein; Modesto Milling, CA), grit and water. Each aviary houses 10-12 breeding pairs of rock doves and includes wooden nest boxes and nesting material (straw). Birds are allowed to naturally form breeding pairs and select and defend nest sites. Nest boxes are checked daily and the identity of the attending parent, presence and number of eggs or chicks is recorded. This daily data collection allowed us to collect samples at precise stages of incubation and yielded a full breeding history for each individual bird.

To examine the effect of prior parental experience on hormonal stress responses in a non-parental state, we also collected blood samples from birds that had (“experienced”), and had not (“inexperienced”), raised chicks in previous nests. Experienced birds had raised at least one chick in a prior nest. Sample sizes were 16 inexperienced birds (8 females, 8 males) and 19 experienced birds (10 females, 9 males). Samples were collected when birds had no active nest. Birds with no active nest can be considered in a non-parental, “baseline” state, as they have no eggs nor chicks to attend, though they are likely reproductively active and may be engaged in courtship or nest building. The average time since the last nest effort was completed did not significantly differ between experienced and inexperienced birds (8.4 days vs. 21.5 days on average; *t* = 1.03, *p* = 0.317). Experienced birds were older than inexperienced birds at sampling time (1.84 years vs. 1.38 years on average; *t* = 3.31, *p* = 0.002). We continued to collect breeding data on these birds after blood samples were collected, and experienced birds initiated a new nest effort (defined as the first day an egg was laid) significantly sooner than inexperienced birds (8.6 days vs. 24.9 days on average; *t* = -1.03, *p* = 0.032).

All methods and procedures were approved by the University of California Davis IACUC (protocol #22407).

#### Dexamethasone dosage validation

To test birds’ maximal negative feedback ability after a stressor, we used dexamethasone (DEX), a synthetic glucocorticoid that selectively binds glucocorticoid receptors to initiate negative feedback and downregulate CORT release (Lattin and Kelly, 2020), including in rock doves (Westerhof et al., 1994). To ensure DEX reduced CORT levels significantly below stress-induced levels and to levels similar to baseline, we conducted a validation experiment with multiple dosages. We captured non-breeding rock doves (total *n* = 19) and placed them in an opaque cloth bag for 30 minutes to simulate an acute stressor. This capture-restraint method is a classic handling stress paradigm that has been used to reliably increase CORT levels in birds (Romero and Wingfield, 2016; Wingfield et al., 1982), including in our rock dove population (Calisi et al., 2018). After 30 minutes, we removed birds from bags, took a ∼100 µL blood sample from the alar wing vein. We then immediately injected birds intramuscularly with DEX (Cat No. D1756, Sigma Aldrich, Milwaukee, WI, USA) at either 1 mg/kg (*n* = 3), 2 mg/kg (*n* = 5), or 4 mg/kg (*n* = 5) or with 0.9% physiological saline as a vehicle control (*n* = 6). Birds were returned to their home cage to recover, then recaptured and bled after an additional 60 and 90 minutes post-DEX (∼ 100 µL each sample). All blood samples were taken between 0800 - 1100 (PST) in February 2020.

Plasma CORT levels in birds treated with saline vehicle did not change after 60 or 90 minutes of recovery post-stressor (Fig.2; one-way ANOVA: *F*_2,16_ = 0.4, *p* = 0.957). Despite being effective in other bird species (M. J. Dickens et al., 2009; Lattin et al., 2012), 1 mg/kg DEX also did not significantly reduce CORT levels 60 or 90 minutes after stress (*F*_2,5_ = 1.1, *p* = 0.396). DEX dosages of 2 mg/kg (*F*_2,12_ = 7.4, *p* = 0.008) and 4 mg/kg (*F*_2,12_ = 7.7, *p* = 0.007) significantly decreased after 60 and 90 minutes recovery. Additionally, both 2 mg/kg and 4 mg/kg DEX doses significantly differed from vehicle after 60 and 90 additional minutes of recovery (*F*_3,16_ = 4.1, *p* = 0.023). Thus, we chose to use the lowest effective DEX dose, 2 mg/kg, measuring post-DEX CORT levels after 60 minutes of recovery.

**Figure 1.**
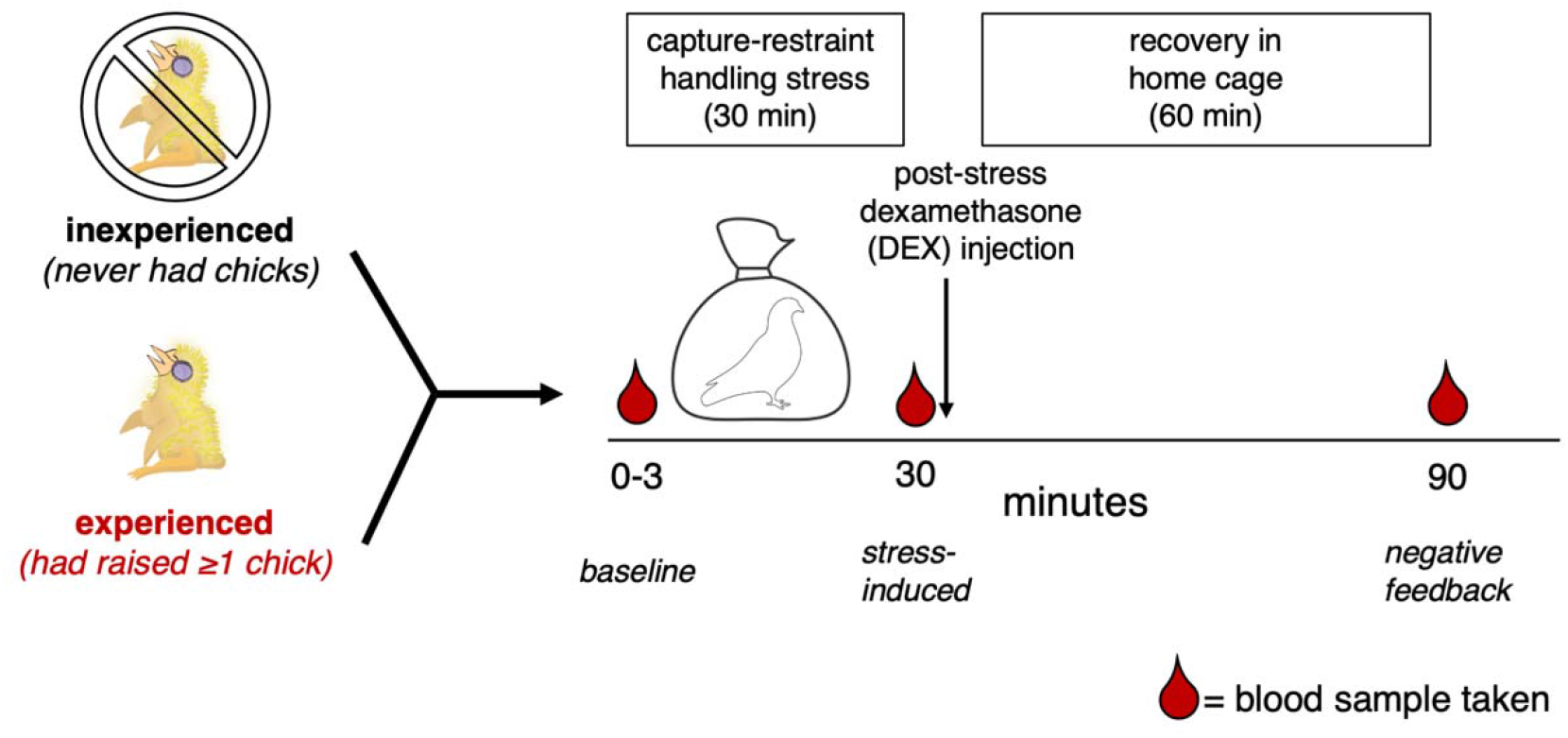
Sampling paradigm for Experiment 1. We collected stress series from inexperienced (never raised chicks, had laid eggs) and experienced birds (had raised at least one chick) that currently had no active nest to understand the influence of prior parental experience on plasma hormone levels after a stressor. Three blood samples were taken from birds to assess hormonal responses to stress: 1) baseline (< 3 minutes from capture), 2) stress-induced (after 30 minutes in an opaque cloth bag, representing a classic capture-restraint stressor), and 3) negative feedback (60 minutes after injection with dexamethasone and recovery in home cage). Dexamethasone was injected immediately after the stress-induced blood sample was taken. Plasma from blood samples were used to measure corticosterone (CORT) and prolactin. Sample sizes for each group can be found in the Methods.

**Fig 2.**
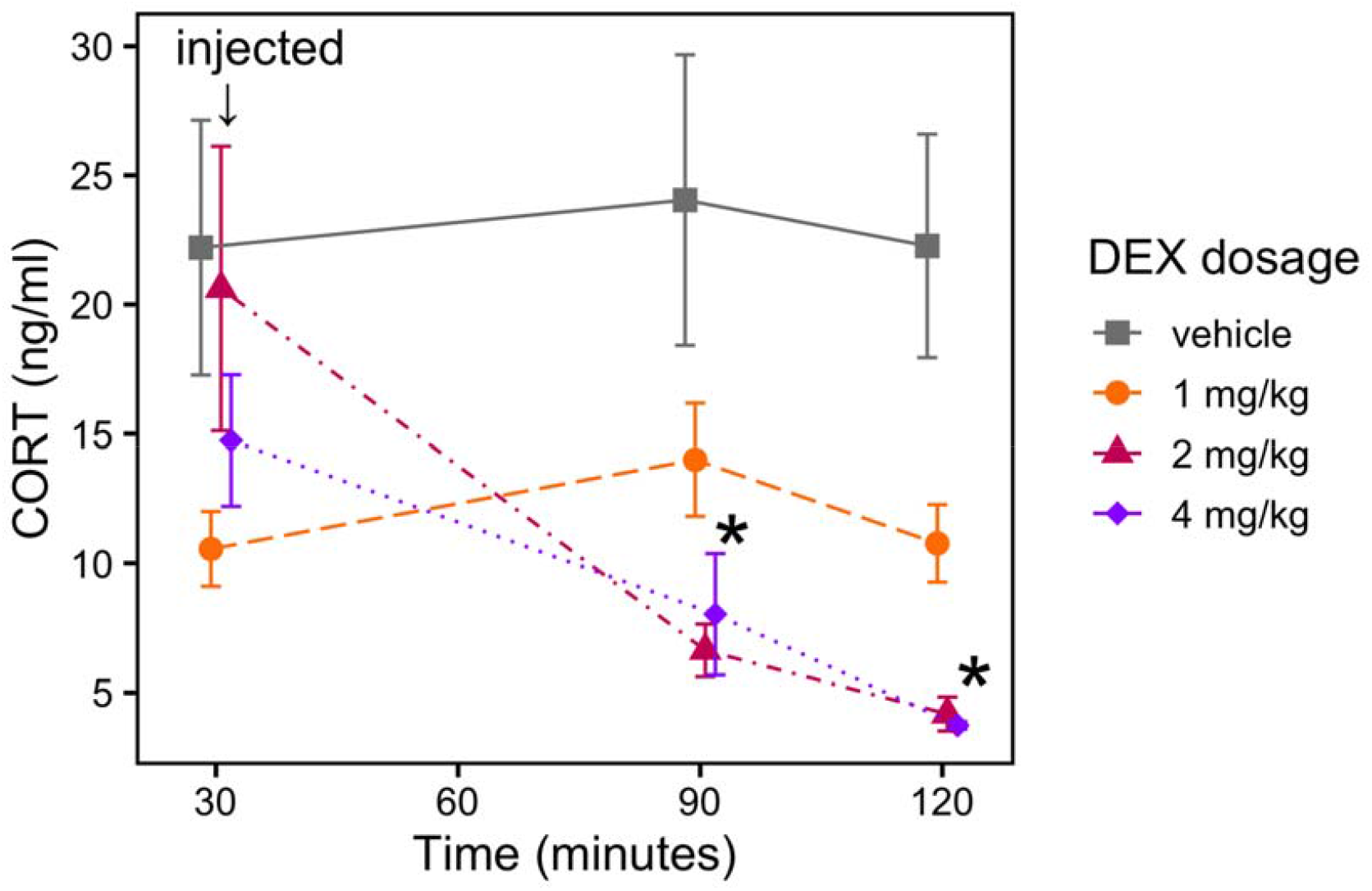
Plasma corticosterone (CORT) response to injections of various doses of dexamethasone (DEX) or vehicle after 30 minutes of restraint. Birds were either injected with a dose of DEX (1 mg/kg, *n* = 3; 2 mg/kg, *n* = 5, 4 mg/kg, *n* = 5) or saline vehicle (*n* = 6) after being exposed to an acute capture-restraint stressor for 30 minutes (injection time indicated with an arrow). Mean ± SEM are shown. ∗= *p* < 0.05 for 2 mg/kg and 4 mg/kg doses compared to the 1 mg/kg and vehicle groups at the same timepoint.

#### Stress series blood collection

For experimental stress series, we collected three blood samples from each bird under the classic capture-restraint protocol (Fig.1A). First, we collected a sample of blood from the alar wing vein using a 26G needle within three minutes of capture (109 ± 36 seconds) from the bird’s home cage. Samples collected within three minutes of capture are considered representative of baseline levels for both circulating CORT and PRL (Chastel et al., 2005; Romero and Reed, 2005), and we found no effect of time to sample on either baseline concentration of either hormone (CORT: *F*_1,28_ = 0.57, *p* = 0.458; PRL: *F*_1,27_ = 0.31, *p* = 0.583). We then placed each bird in an opaque cloth bag to simulate an acute handling stressor and collected a second blood sample 30 minutes later to measure stress-induced hormone levels. After this blood sample was taken, we injected each bird intramuscularly with 2 mg/kg DEX (dosage validated as described above) and then returned birds to their home cage to recover. We collected the last blood sample 60 minutes after DEX injection to measure negative feedback hormone levels. All blood samples were approximately 100 µL each and were collected between 0800 - 1200 PST in April - June 2021. We found no effect of time of day on either CORT (*F*_1,100_ = 0.01, *p* = 0.939) or PRL concentration (*F*_1,97_ = 0.19, *p* = 0.660).

We centrifuged blood samples for 10 minutes at 10,000 rpm to separate plasma. Plasma aliquots were then stored at -80□ until further analysis. All samples were brought up to 4□before being run in immunoassays.

#### Corticosterone (CORT) radioimmunoassays

Circulating corticosterone (CORT) concentrations were measured from plasma at the UC Davis Metabolomics core using a commercially available radioimmunoassay (RIA) kit (MP Biomedicals, Orangeburg, NY). A serial dilution was performed prior to the assay, and plasma samples from 0 min, 30 min and 90 min timepoints were run at 1:11, 1:26, and 1:11 dilutions, respectively. Cross-reactivity with *C*.*livia* CORT was validated previously for this assay (Austin et al., 2021a; Calisi et al., 2018), and the assay had a limit of detection of 0.0385 ng/ml.

Samples were run in duplicate, and mean intra-assay and inter-assay coefficients of variation (CV) were 5.0% and 6.5%, respectively. All samples from the DEX dosage validation were run in a single assay.

#### Avian prolactin enzyme-linked immunoassay (ELISA)

We measured circulating prolactin (PRL) using a heterologous competitive enzyme-linked immunoassay (ELISA) using the methods described in detail in (Smiley and Adkins-Regan, 2016) and developed by ADS Biosystems, Inc. (San Diego, CA). This assay has previously been used with rock dove plasma (Booth et al., submitted). The full protocol can be accessed at: protocols.io: dx.doi.org/10.17504/protocols.io.36wgq7zoovk5/v1.

Briefly, biotinylated recombinant chicken PRL (ADS Biosystems, Inc., San Diego, CA) is added to samples and standards and competes for binding sites on the bound rabbit anti-chicken PRL antibodies (A.F. Parlow, National Hormone and Peptide Program, Los Angeles, CA). Visualization occurs through an enzymatic reaction using streptavidin horseradish peroxidase (HRP) (Cat. No. 21130, Thermo Fisher, Waltham, MA). Chicken PRL antibodies have been successfully used in ELISA to measure prolactin from other avian species, including zebra finches and brown-headed cowbirds (Lynch et al., 2020; Smiley and Adkins-Regan, 2016). We confirmed parallelism of serially-diluted *C*.*livia* plasma with a chicken PRL standard curve, and spike-recovery of chicken PRL spiked into a *C*.*livia* plasma sample. We used two pooled validation samples, one from non-breeding birds (low pool) and one from birds at incubation day 17 (high pool) to calculate intra-assay CV. Mean intra-assay CV was 5.79%. We ran all plasma series (0 min, 30 min, and 90 min samples) for an individual bird on the same ELISA plate. All samples were run in duplicate, along with a standard curve on each 96-well plate. Plates were read on an iMark microplate reader (Bio-Rad Laboratories, Hercules, CA) at 450 nm with background subtracted from 595 nm. Concentrations were interpolated from the standard curve using a 4-parameter fit (iMark software v.1.04, Bio-Rad Laboratories). One individual (inexperienced no-active-nest) was not run due to hemolysis that contaminated the plasma samples.

#### Statistical analysis

All statistical analyses were performed in the R statistical language (v.4.0.3). For each hormone (CORT and PRL), we created a mixed effects linear model, where a random effect of individual bird was included to account for the repeated-measures design of our stress series. Models were created using the lme4 package (v.1.1.27)(Bates et al., 2015). In this model, we tested how the independent variables of experience level (experienced with chicks or inexperienced), stress series timepoint, sex, and their interactions affected the dependent variable of hormone concentration. To improve distribution of the data, all hormone concentrations were log_10_-transformed. We report results from ANOVA (Type III) run on these mixed effects models using the car package (v.3.0.1)(Fox and Weisberg, 2019). We also report the results of post-hoc comparisons performed with estimated marginal means, corrected with Benjamini-Hochberg false discovery rate (FDR) corrections in the emmeans package (v.1.5.2)(Lenth, 2020). All code for statistical analysis and figures can be found at: https://github.com/vsfarrar/experience-stress-hippocampus.

### Experiment 2: Hippocampal and pituitary gene expression

In a second experiment, we extended results from Experiment 1 that showed an effect of parental experience on CORT and PRL release after an acute stressor. Here, we examined gene expression in brain and pituitary tissues collected from birds with and without prior experience with chicks, to determine if genes involved in stress response regulation were differentially expressed in experienced birds versus inexperienced ones. Specifically, we examined glucocorticoid receptor (GR; also known as NR3C1) and mineralocorticoid receptor (MR, or NR3C2) in the hippocampus, as these two receptors are known to regulate negative feedback of the glucocorticoid stress response and HPA axis regulation (Herman et al., 2012; Phillips et al., 2006).

#### Tissue collection

Whole brains were collected from 30 reproductively mature rock doves (age range: 1 -2 years old) that were not actively nesting. Of these, 16 (n = 8 males, 8 females) birds had previously raised at least one chick (average chicks raised: 2.5 ± 1.46), and 14 (n = 7 males, 7 females) had never raised chicks. The mean time since birds last had an active nest (at time of collection) did not significantly differ with experience (6.6 days for experienced vs. 10.0 days in inexperienced birds: *t* = 0.85, *p* = 0.419). As in above, experienced birds were older than inexperienced birds, but only by a matter of days; all birds were between one and two years old (average age: 1.41 years for inexperienced vs 1.66 years for experienced individuals, *t* = -2.16, *p* = 0.041).

We euthanized birds using methods previously used in rock doves (Calisi et al., 2018; MacManes et al., 2017). Within three minutes of capture, birds were euthanized with an overdose of isoflurane then rapidly decapitated. Whole brains and pituitary glands were removed and flash-frozen on dry ice, then stored at -80□until further analysis. All tissues were collected between 0800-1100 PST in March 2020. As these methods are terminal, different individual animals were used in this experiment than those in Experiment 1.

#### Hippocampi microdissection

To analyze gene expression specific to the hippocampus, we microdissected the hippocampus from whole brains using a Leica CM1950 cryostat. We collected hippocampus tissue using a 3 mm diameter punch from 100 µM slices. We used landmarks from the Karten and Hodos (1966) pigeon brain atlas to locate the hippocampus, starting with when the commissura anterior (CA) visibly crossed the coronal section and ending when the cerebellum was visible (Fig.4A, plates A 7.75 - A 4.25 in Karten and Hodos 1966; average of 27-30 punches at 100 µM). Hippocampus tissue punches were stored in 200 µL TriSure Reagent (BioLine, Meridian Bioscience, Cincinnati, OH, USA) at -80□until RNA extraction.

#### Quantitative PCR

To extract total RNA from hippocampal tissue, we used a modified protocol of the Direct-zol RNA extraction kit (Catalog No. R2501, Zymo Scientific, Irvine, CA, USA) along with TriSure reagent (Catalog No. 38032, BioLine, Meridian Bioscience). A full RNA extraction protocol can be accessed online at protocols.io: dx.doi.org/10.17504/protocols.io.5qpvob6p9l4o/v1.

RNA concentration and quality was measured using a Nanodrop ND-1000 spectrophotometer (Thermo Fisher Scientific, Waltham, MA, USA). All samples passed quality assurance and had 260/280 ratios and 260/230 ratios > 1.80. We removed any remaining genomic DNA from RNA samples using Quanta Perfecta DNase I (Catalog No. 95150-01K, Quanta Biosciences, Gaithersburg, MD, USA). We then converted RNA to single-stranded complementary DNA (cDNA) via reverse transcription using qScript cDNA Supermix (Catalog No. 95048-100, Quanta Biosciences). We diluted total cDNA 5-fold in preparation for qPCR.

Using quantitative PCR (qPCR), we measured relative gene expression for glucocorticoid receptors (*GR*) and mineralocorticoid receptors (*MR*) in the hippocampus. We also measured expression of two reference genes, *HPRT1*, and *RPL4*, to account for variation in total transcription between samples. All primers were designed using the *Columba livia* transcriptome v2.10 (NCBI accession no. GCA_000337935.2) as a template. We also validated each primer for ideal replication efficiencies and singular melt curves using a standard curve consisting of five 10-fold dilutions of pooled tissue cDNA. Primer details can be found in Table 1.

**Table 1.**
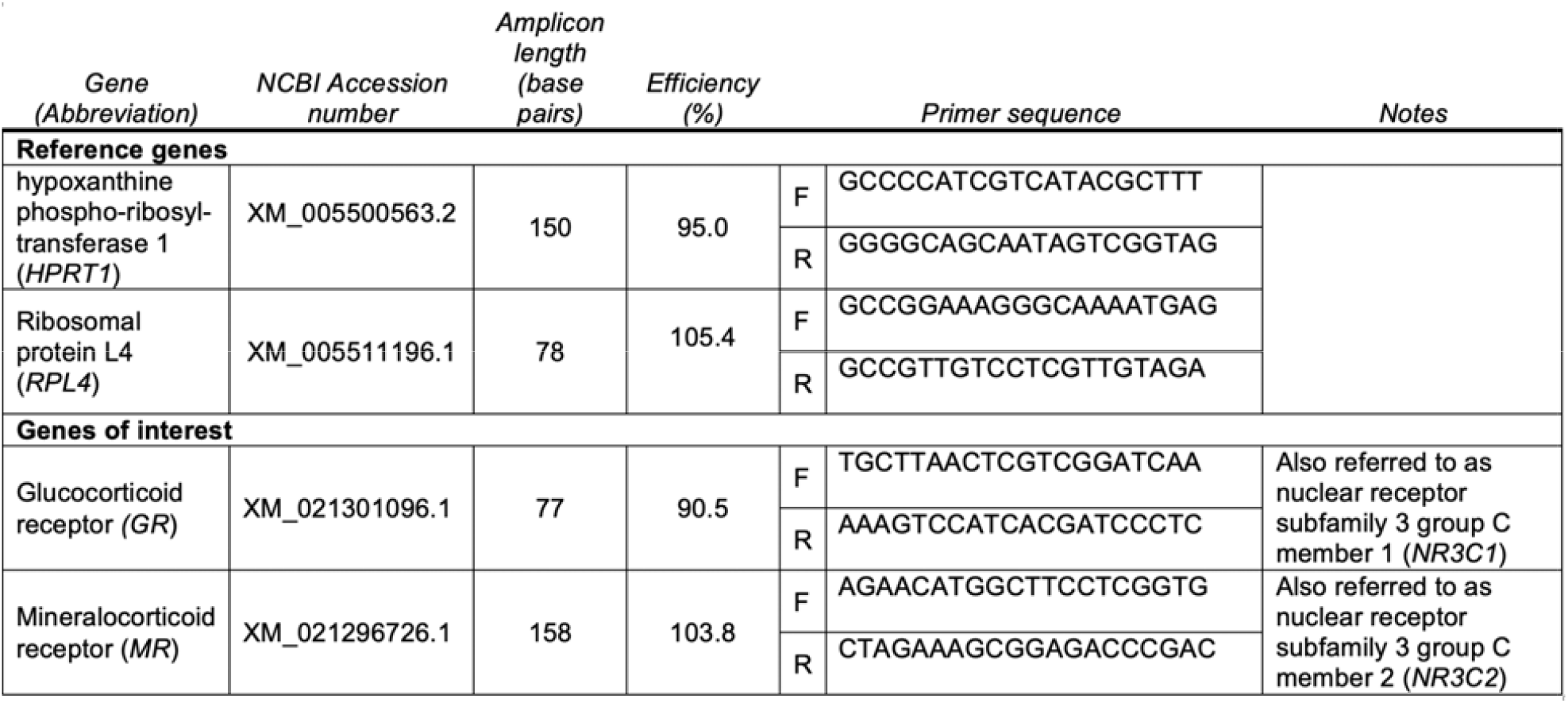
Primers used in quantitative PCR. Primers were designed using the NCBI Primer-BLAST tool on gene templates for *Columba livia* (indicated by NCBI accession numbers). Primer efficiencies were calculated by running a standard curve of five 10-fold dilutions of purified PCR product, and primers were evaluated for single products via melt-curve analysis during validation.

We ran each sample in triplicate on a 384-well plate using the following qPCR reaction mix: 1 µL diluted cDNA template, 5 µL 2X SSOAdvanced SYBR Green PCR mix (BioRad), and 10 µM each of primer (total volume: 10 µL, Invitrogen). We ran plates on a CFX384 Touch Real-Time PCR detection system (BioRad) under the following thermocycling protocol: 50□for 2 min, 95°C for 10 min, and then 40 cycles of 95□for 15 sec and 60°C for 30 sec. Plates also included no-template controls. We ran all samples of each tissue-gene combination on a single plate.

To calculate relative gene expression from raw qPCR data, we used the Livak and Schmittgen (2001) delta-delta Ct method. To do this, we first normalized the expression (cycle threshold, Ct) of each gene of interest to the geometric mean of reference gene expression for that sample. We used *HPRT1* and *RPL4* as reference genes, as recommended for avian neural tissue (Zinzow-Kramer et al., 2014). We verified that expression of these reference genes did not differ with parental experience (*F*_1,27_ = 2.14, *p* = 0.155) or sex (*F*_1,27_ = 0.09, *p* = 0.766) in our samples using two-way ANOVA. Then, we relativized normalized expression (delta-Ct) for each sample to the average normalized expression for the control group (delta-delta-Ct), which in this case was inexperienced birds. Lastly, we calculated fold change, or 2^(-ddCt)^. Fold change was log_10_-transformed for statistical analyses.

#### Statistical analysis

For each gene of interest (*GR* or *MR*), we ran a linear model to test how the dependent variable, log fold change, may be affected by the independent variables of experience with chicks, sex, and their interaction. We also calculated the ratio of *MR t*o *GR* expression (MR:GR ratio) and examined whether this ratio was also affected by experience with chicks, sex, or their interaction using a linear model. We report ANOVA based upon these linear models.

We ran each gene in a separate model because 1) different transcription factors and promoters are known to underlie expression of these receptors (Biddie and Hager, 2009; Herman and Spencer, 1998) and 2) direct comparisons are not recommended due to the relative expression calculations used in the Livak and Schmitgen (2001) method.

## Results

### Experiment 1: Hormonal responses to acute stress

When compared between birds that were not actively nesting (i.e., not actively caring for eggs or chicks in nests), previous parental experience with chicks significantly altered the CORT and PRL stress responses. We found a significant interaction between experience and stress-series timepoint on CORT levels (Table 2). Post-hoc analyses show that this interaction is driven by experienced birds having lower CORT post-stressor (*t* = -2.18, *p* = 0.033) and after DEX-induced negative feedback (*t* = -2.63, *p* = 0.011), but not at baseline (*t* = -0.19, *p* = 0.853) (Fig.3A). Further, timepoint as a main effect was significant (Table 2), as expected. Averaged over levels of experience and sex, CORT levels increased in response to 30 minutes of acute-restraint stress compared to baseline (*t* = -23.5, *p* < 0.001) and subsequently decreased after 60 minutes of DEX-induced negative feedback (*t* = 14.6, *p* < 0.001). Negative feedback CORT levels were also significantly higher than baseline levels (*t* = -9.14, *p* < 0.001). There was no significant main effect of sex, nor a significant interaction between sex and experience or sex and timepoint in CORT levels (Table 2).

**Fig 3.**
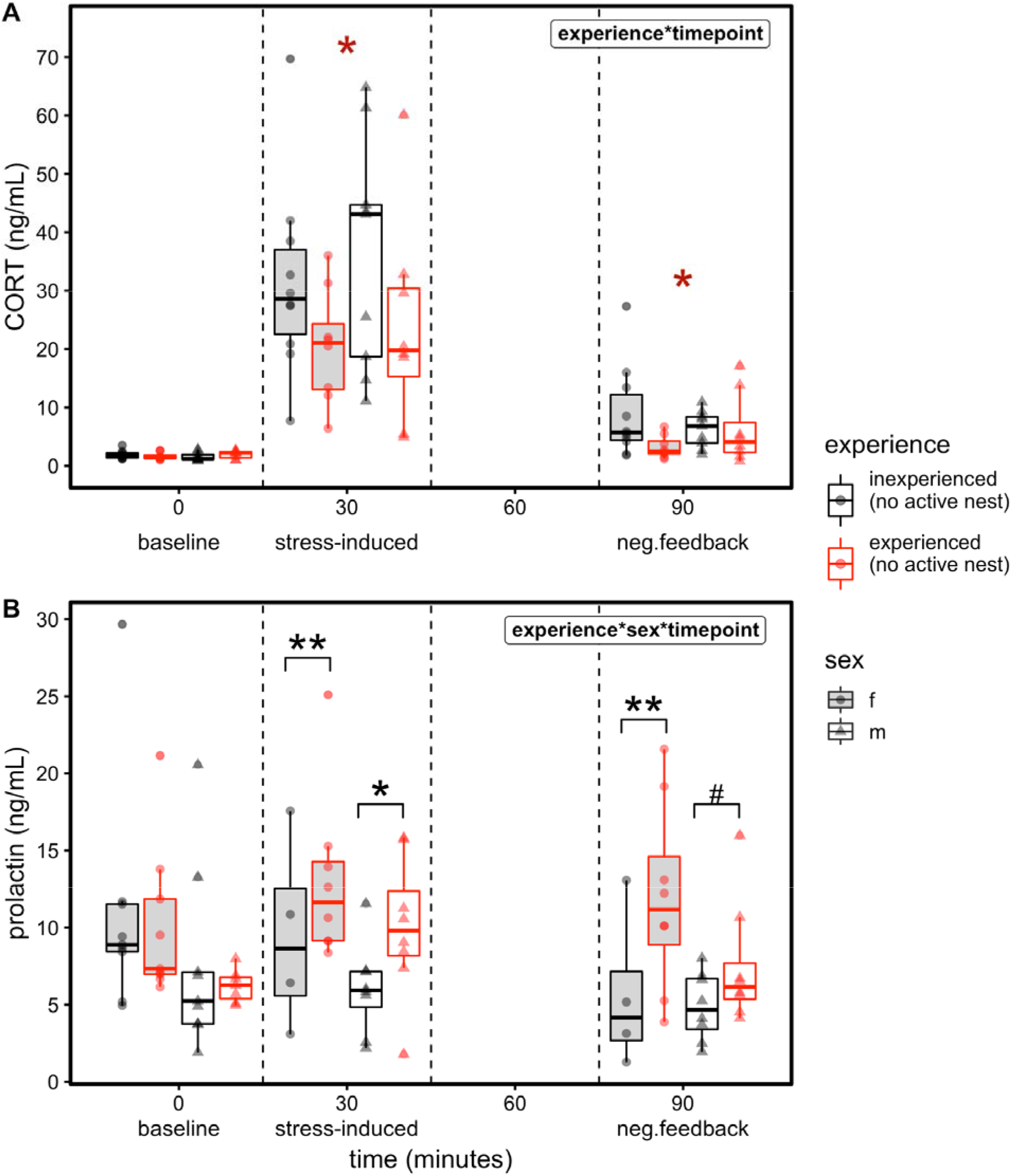
Circulating hormones vary with parental experience, stress-series timepoint, and sex. (**A**) Plasma corticosterone (CORT) and (**B**) plasma prolactin was measured in birds withou active nests that varied in previous parental experience with chicks (coded by line color; black and red represent inexperienced and experienced birds, respectively). Hormones were measured at baseline (0-3 minutes after capture), after capture-restraint stress (30 min post capture) and after dexamethasone (DEX) induced negative feedback (60 minutes of recovery post stressor, 90 min after capture). Sampling timepoints are separated visually with dashed lines. Points represent individual birds, and boxplots represent the where the first quartile, median, and third quartile for each sex and stage. These stress series were collected for both females (circles; boxplots shaded in gray) and males (triangles, boxplots unshaded). The highest-level, significant predictors from the linear mixed model that included experience, timepoint, sex and their interactions are shown in bold in the upper right corner (see Table 2). In plot A, red asterisks (*) indicate a significant effect (*p* < 0.05) of experience averaged over levels of sex in post-hoc analyses. In plot B, brackets represent significant differences in experience between the sexes across timepoints in post-hoc contrasts, where # = *p* < 0.1, * = *p* < 0.05, and ** = *p* < 0.01.

**Table 2.** ANOVA from mixed effect model for effects of parental experience, stress series timepoint, sex and their interactions on log_10_-transformed concentrations of corticosterone and prolactin. This model included a random effect of individual bird to account for the repeated measures design of the stress series. All birds included in this dataset were not actively nesting (i.e., not actively incubating eggs or caring for chicks). Significant effects at the □= 0.05 level are indicated in bold.

| Variable | Corticosterone |  |  | Prolactin |  |  |
| --- | --- | --- | --- | --- | --- | --- |
|  | df | F | p | df | F | p |
| Experience | 1 | 4.24 | **0.040** | 1 | 16.49 | **<0.001** |
| Sex | 1 | 0.01 | 0.913 | 1 | 0.04 | 0.844 |
| Timepoint | 1 | 568.07 | **<0.001** | 1 | 12.29 | **0.002** |
| Experience x Sex | 2 | 0.39 | 0.532 | 2 | 3.02 | 0.082 |
| Experience x Timepoint | 1 | 6.30 | **0.043** | 1 | 29.88 | **<0.001** |
| Sex x Timepoint | 2 | 0.91 | 0.634 | 2 | 5.05 | 0.080 |
| Experience x Sex x Timepoint | 2 | 2.07 | 0.357 | 2 | 7.30 | **0.026** |
| Residuals | 92 |  |  | 89 |  |  |

In PRL levels, we found a significant three-way interaction between experience, sex, and timepoint (Table 2). This interaction implies that previous parental experience affects the stress response sequence differently between the sexes. As with CORT, we found that experience altered levels across timepoints when averaged across sexes (Table 2), with experienced birds having higher levels of PRL both 30 (*t* = 4.85, *p* < 0.001) and 90 minutes (*t* = 4.99, *p* < 0.001) after a stressor. However, this relationship differed significantly between the sexes. The difference between experienced and inexperienced females was larger at both 30 and 90 minutes than the difference between experience levels in males at these timepoints (Fig.3B; Supplemental Figure 1). At 30 minutes, experienced females had 4.9 ± 1.6 (SE) times higher PRL than inexperienced females (*t* = 4.84, *p* < 0.001), while experienced males only had 1.9 ± 0.6 times higher PRL than inexperienced males (*t* = 2.02, *p* = 0.048). Similarly, after 90 minutes, experienced females had 5.8 ± 1.8 times higher PRL levels compared to their inexperienced counterparts (*t* = 5.24, *p* <0.001), compared to only 1.8 ± 0.6 times for males (*t* = 1.82, *p* = 0.073) (Fig.3B). Overall, the shape of the PRL stress response differed with experience when birds were not actively nesting, with experienced birds showing a slight, but not significant, PRL increase post-stressor, and inexperienced birds showing the typical, significant decrease after an acute stressor (Fig.3B; Supplemental Figure 1).

#### Experiment 2 : Hippocampal and pituitary gene expression

In the hippocampus, prior experience with chicks significantly increased *GR* gene expression (Fig.4B; *F*_1,26_ = 11.1, *p* = 0.002) but did not affect *MR* gene expression (Fig.4C; *F*_1,26_ = 2.7, *p* = 0.113). There was no significant effect of sex (*F*_1,26_ = 0.5, *p* = 0.530) nor a significant interaction between sex and parental experience (*F*_1,26_ = 2.0, *p* = 0.164) on *GR* expression. However, *MR* expression did show a significant interaction between sex and experience with chicks (*F*_1,26_ = 6.7, *p* = 0.015). This effect appears to be due to inexperienced females having significantly lower *MR* expression than inexperienced males (*t* = -2.86, *p* = 0.007), but this sex difference in *MR* expression is not present in experienced birds (*t* = 0.71, *p* = 0.486). However, we found no significant effect of experience (*F*_1,26_ = 0.9, *p* = 0.350), sex (*F*_1,26_ = 2.5, *p* = 0.127), nor their interaction (*F*_1,26_ = 2.6, *p* = 0.120) on MR:GR expression ratio.

**Figure 5.**
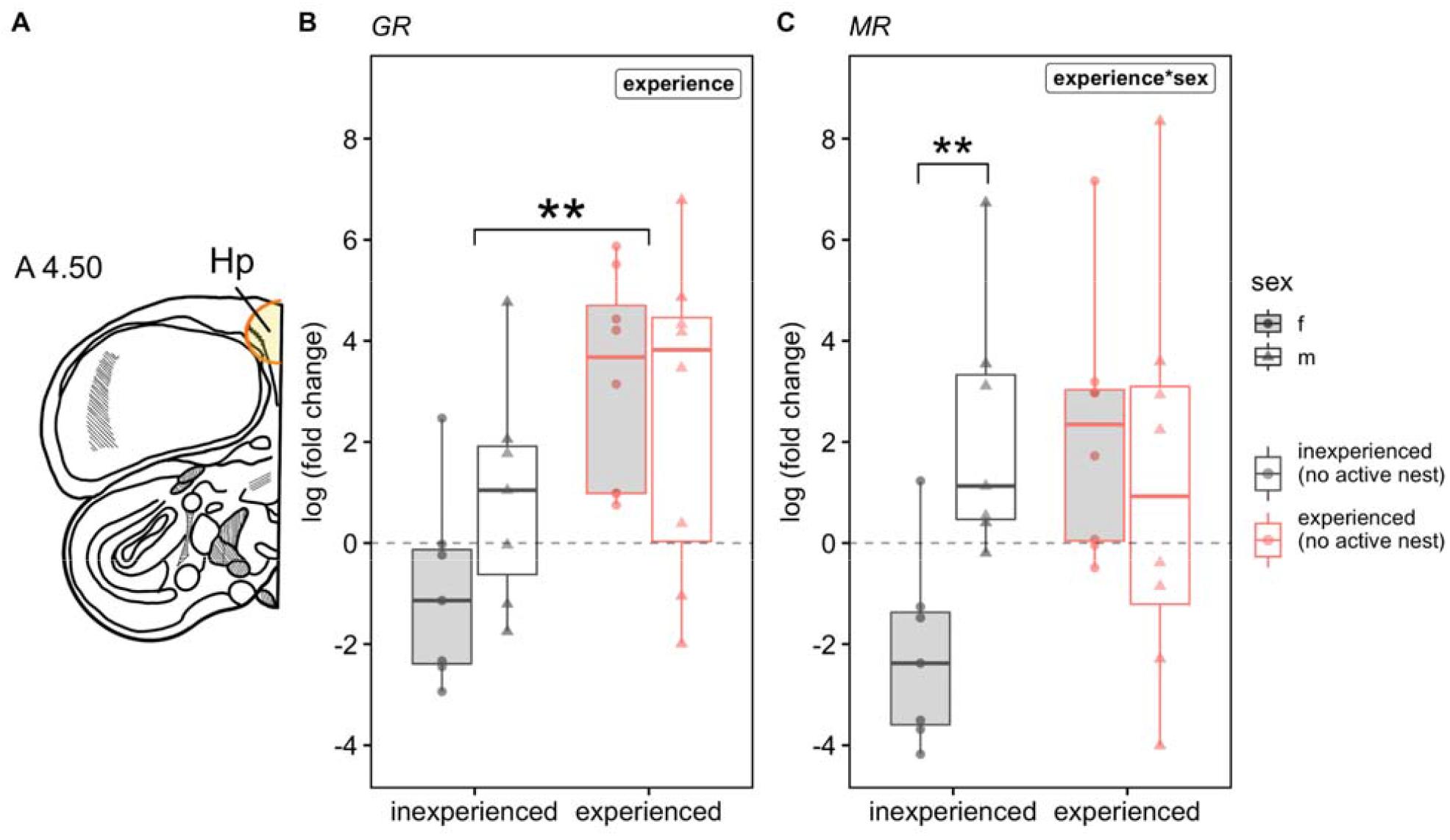
Relative expression of glucocorticoid receptor types in the hippocampus of birds with and without prior parental experience with chicks. **(A)** Representative hippocampal sections, in which (B) glucocorticoid and (C) mineralocorticoid receptor expression was measured using quantitative PCR and compared across birds who had never previously raised chicks (“inexperienced”, gray) and birds who had previously raised at least one chick (“experienced”, red). Points represent individual birds, and boxplots represent the where the first quartile, median, and third quartile for each sex and experience level. Sex is denoted by boxplot fill and point shape (females with shaded boxplots, circles, and males with unshaded boxplots, triangles). Significant predictors from the linear model including experience, sex and their interactions are shown in bold in the upper right corner (see Results). Brackets indicate specific significant post-hoc comparisons after FDR correction, with ** = *p* < 0.01.

## Discussion

We found that previous parental experience with chicks decreased stress-induced and dexamethasone-induced negative-feedback CORT levels and led to increased stress-induced PRL in rock doves without active nests (i.e., in a pre-parental state). Further, in a separate experiment, we found that birds of both sexes with previous experience with chicks also had higher hippocampal *GR* than inexperienced birds. Increased *GR* may lower the threshold for negative feedback and suppressive effects on the HPA axis in experienced birds (Sapolsky et al., 2000), thus potentially mediating the changes in the CORT stress response we observed. Together, these results suggest that inexperienced birds may be constrained by their HPA axis physiology and may not be able to attenuate their stress responses to prioritize future reproduction (support for the “constraint” hypothesis; Curio 1983).

### Effects of parental experience on hormonal stress responses

Prior parental experience with chicks led to lower CORT, and higher PRL levels, both after an acute stressor and after dexamethasone-induced negative feedback. Previous studies that examined effects of prior breeding experience on CORT and PRL only measured baseline hormone levels (Angelier et al., 2007b, 2006), and found that experienced albatross had higher baseline CORT and PRL during brooding than birds breeding for the first time. Higher baseline PRL has also been found in experienced zebra finches (Smiley and Adkins-Regan, 2016) and cotton-top tamarin monkeys (Ziegler et al., 1996) during breeding. However, we did not find any significant effects of experience on baseline CORT or PRL levels in pre-parental birds with no active nest. Experience only significantly altered hormone levels after a stressor or during negative feedback in our study, highlighting the importance of measuring hormone responses beyond baseline levels to understand HPA axis plasticity. Although our findings did not align with previous work on breeding experience, they did mirror patterns seen with increasing age. In common terns, a long-lived seabird, CORT and PRL were lower and higher, respectively, after acute restraint stress in older parents compared to younger ones during incubation (Heidinger et al., 2010, 2006). Similarly, younger snow petrel females had lower stress-induced PRL than older females (Angelier et al., 2007a), and senescent albatross had lower CORT levels, but not PRL, levels than younger birds (Angelier et al., 2006). In our study, the range of ages was small (0.5 - 3 years, with 80% between 1-2 years old), making age less likely to drive our observed effect of experience. Instead, increasing age likely correlates with breeding experience in other study populations, so differences in stress responses with age seen in prior work may be mediated in part by parental experience. Indeed, when both were measured, breeding experience appeared to better statistically predict hormone levels than age (Angelier et al., 2007b, 2006).

Our observation that birds without prior parental experience exhibit a more reactive stress response in both CORT and PRL than experienced birds lends support for both the “constraint” and “restraint” hypotheses about why reproduction may improve with age (Curio 1983). Under the constraint hypothesis, inexperienced birds may be limited (constrained) in their ability to invest in reproductive efforts over personal survival in the face of stressors. That is, inexperienced birds may not be able to modulate down and attenuate the HPA axis or maintain PRL secretion under stress. This interpretation implies that there may be mechanistic differences in HPA regulation between inexperienced and experienced birds, which we found evidence for in the hippocampus (see below). Alternatively, as the inexperienced birds we sampled were slightly, but significantly, younger than experienced birds (mean 1.84 vs 1.38 years), the “restraint” hypothesis may also be supported (Curio 1983). In this case, inexperienced, younger birds may limit (restrain) their parental investment due to their relatively larger opportunities for future reproduction compared to older, more experienced breeders (Stearns, 1976). This interpretation is also consistent with the “brood value hypothesis” (Heidinger et al., 2010; Lendvai et al., 2007), where older, experienced birds may modulate their stress response because their current / next brood has relatively higher reproductive value.

Another interpretation is that experienced birds were closer to a parental state than inexperienced birds, driving stress response differences. This explanation would occur if experienced birds had more recently ended a chick care bout or were closer to restarting their next nest than inexperienced birds. Although birds did not have active nests when sampled, experienced birds did initiate new nests sooner after sampling than inexperienced birds on average (8.6 vs 24.9 days), though the time since last nest effort did not differ significantly (9.9 vs 14.9 days). Thus, we cannot rule out that the effects of experience may be due to differences in reproductive state or engagement in pre-parental behaviors. Even under this interpretation, however, our results would still be consistent with the “parental care hypothesis” (Wingfield et al., 1995). This hypothesis states that birds more involved in parental effort show attenuated stress responses than those not engaged in care. Comparing stress responses in birds of varying experience during the parental period (i.e., during incubation or brooding) would clarify if our results are due to differential reproductive states or truly represent a persistent effect of experience.

### Effects of parental experience on hippocampal glucocorticoid receptors

When we examined hippocampal glucocorticoid receptors, we found that, when not actively nesting, birds of both sexes that had previously had chicks had higher *GR* expression than birds inexperienced with chicks. Combined with our hormonal stress response results, this suggests that increased hippocampal *GR* may allow experienced birds to enact negative feedback on their HPA axis more rapidly and/or at a lower threshold level of circulating CORT, leading to overall lower stress-induced and negative-feedback CORT compared to inexperienced birds. Thus, hippocampal receptors provide a potential molecular mechanism for the “constraint” hypothesis, where young, inexperienced birds may be limited (constrained) in their ability to attenuate stress responses and prioritize current reproductive efforts (Curio, 1983). While modified hippocampal GR expression has been a target of interest of the effects of stress in early development (Harris and Seckl, 2011; Lupien et al., 2009), our finding opens further investigation into plasticity of this mechanism in adults. However, it remains unclear whether the differences with experience we observed persist throughout the parental care period, which would be important to establish in future studies.

Our results contrast with previous work in avian species, which suggests that the hippocampal MR may be more important in modulating the glucocorticoid stress response than GR. For example, hippocampal MR expression, but not GR, was altered in zebra finch lines selected for highly-responsive HPA axes (i.e. high stress-induced CORT) (Hodgson et al., 2007). Developmental stress, such as egg CORT injections or postnatal food restriction, affected hippocampal *MR*, but not *GR*, in Japanese quail (Soleimani et al., 2011; Zimmer and Spencer, 2014). Similarly, chronic stress nor translocation to captivity, which both led to attenuated HPA axis responses, affected hippocampal GR in starlings or chukar (Dickens et al., 2009; Dickens et al., 2011). Alternatively, GR in the hypothalamus, another potential site of negative feedback (Smulders, 2021), may be more important for HPA axis regulation in other species, as chronic and prenatal stress reduced GR in the hypothalamus of European starlings and Japanese quail, respectively (Dickens et al., 2009; Zimmer and Spencer, 2014). Other studies of seasonal transitions, however, found no differences in hippocampal or hypothalamic GR across breeding stages when stress responses had been shown to attenuate (Gambel’s white-crowned sparrows (Krause et al., 2015) ; house sparrows (Lattin and Romero, 2013)). However, our results align more closely with mammalian studies, where changes in hippocampal *GR* affected stress-induced CORT release (Harris et al., 2013; Ratka et al., 1989; van Haarst et al., 1996).

Similarly, we did not find an overall effect of experience on hippocampal *MR* expression, in contrast with other studies that found altered hippocampal *MR* in birds. In the aforementioned studies, selection for highly-reactive stress profiles, chronic stress, developmental stressors, and breeding transitions all altered hippocampal *MR* expression (Dickens et al., 2009; Hodgson et al., 2007; Krause et al., 2015; Zimmer and Spencer, 2014), with all associating decreased *MR* expression with reduced stress-induced CORT release. Again, we did not find this effect. However, we found an apparent sex difference present in inexperienced birds, with females expressing lower *MR* in males, that was not present in experienced birds. While this result suggests that inexperienced females may have lower *MR* densities, allowing *GR* to be bound more rapidly, potentiating faster negative feedback, this was not borne out in the plasma CORT data. Most previous studies only measured these receptors in one sex, though those that included both sexes found no significant differences in both stress response and hippocampal *MR* (Dickens et al., 2009; Hodgson et al., 2007). These results emphasize the importance of studying these mechanisms in both sexes, as to further understand what contexts may lead to presence, and absence, of sex differences in HPA axis regulation.

The discrepancies we found with other avian studies may be due to differences in the context of our study (parental experience) and/or species differences. To our knowledge, our study is the first to investigate how prior parental experiences may affect hippocampal *GR* and *MR* expression in birds. It is possible that prior reproductive cycles, and the many endocrine changes involved (Austin et al., 2021b), may alter hippocampal gene regulation in ways that differ from those observed in stress contexts or other annual cycle transitions. Indeed, female rats show attenuated stress response and reduced hippocampal *GR* expression in late pregnancy (Johnstone et al., 2001). The experience of the changing hormonal milieu during gestation and preparation for lactation may be responsible for some of these changes (as suggested by (Torner and Neumann, 2002). In our study, it remains to be seen if the effects of experience continue beyond the pre-parental stage, when birds enter their next nesting attempt. If these effects truly persist into future breeding efforts, manipulations of hormones involved in the gaining of parental experience, such as fluctuating prolactin or oxytocin/mesotocin, could uncover the causes of HPA axis regulation in the hippocampus. Additionally, negative feedback may be mediated through other mechanisms,, such as steroid-metabolizing enzymes in target cells or corticosterone-binding globulins in plasma (Wingfield et al., 2015). Finally, we cannot rule out the role of species differences, as the continuously-breeding rock dove may differ in stress regulation from the seasonal breeders previously mentioned. Prior work suggests rock doves can regulate CORT differently from other birds in some ways, such as not downregulating HPA activity during molt (Romero and Wingfield, 2001).

## Conclusions

Overall, we found evidence in support of the “constraint” and “restraint” hypotheses for why younger, inexperienced birds may be poorer breeders than older, experienced individuals, that this effect may be related to the ability to attenuate the CORT and PRL stress responses. In turn, the ability of experienced birds to attenuate hormonal stress responses, specifically CORT release, may be mediated by increased hippocampal glucocorticoid receptors involved in HPA axis regulation. We found mixed evidence for the parental care hypothesis and prolactin stress hypothesis across incubation in the rock dove, but future work comparing parental care stages that are distinct in behavior, energetic demands, and offspring cues may further test these hypotheses. Overall, investigations of the effect of parental experience on hormonal stress responses and neural HPA axis regulation are few, and the results here may provide potential mechanisms for further exploration. These results set the stage for future studies examining how experience may enact lasting changes in HPA axis regulation, such as epigenetic mechanisms (Rubenstein et al., 2016; Siller and Rubenstein, 2019), as well as link these mechanisms to behavioral and fitness consequences of gaining parental experience.

## Acknowledgements

James Graham (UC Davis Metabolomics Core) provided invaluable assistance with CORT RIAs. Zhiyong Wang (ADS Biosystems) provided technical support with development and validation of prolactin ELISAs. A.F. Parlow (National Hormone and Peptide Program) graciously provided rabbit anti-chicken PRL antibodies for prolactin ELISAs. We also thank the many undergraduates of the Calisi lab who assisted with animal husbandry and breeding data collection: A. De Matos, D.Erenstein, C. Fargeix, L. Flores, A. Quezada, E. Moges, R. Moore, A. Ramirez, Z. Ricard, J. Robles-Diaz, G. Virata, and B.Wallen.

## CRediT Authorship Statement

**Victoria S. Farrar**: Conceptualization, Methodology, Validation, Investigation, Formal analysis, Writing - Original Draft; **Jaime Morales Gallardo**: Investigation; **Rebecca M. Calisi:** Resources, Supervision, Funding acquisition, Writing - Review & Editing

## Data Availability Statement

Data for this paper can be found on the Dryad data repository at https://doi.org/10.25338/B8KK91. Analysis code can be found at: https://github.com/vsfarrar/experience-stress-hippocampus.

## Supplemental

**Supplemental Figure 1.**
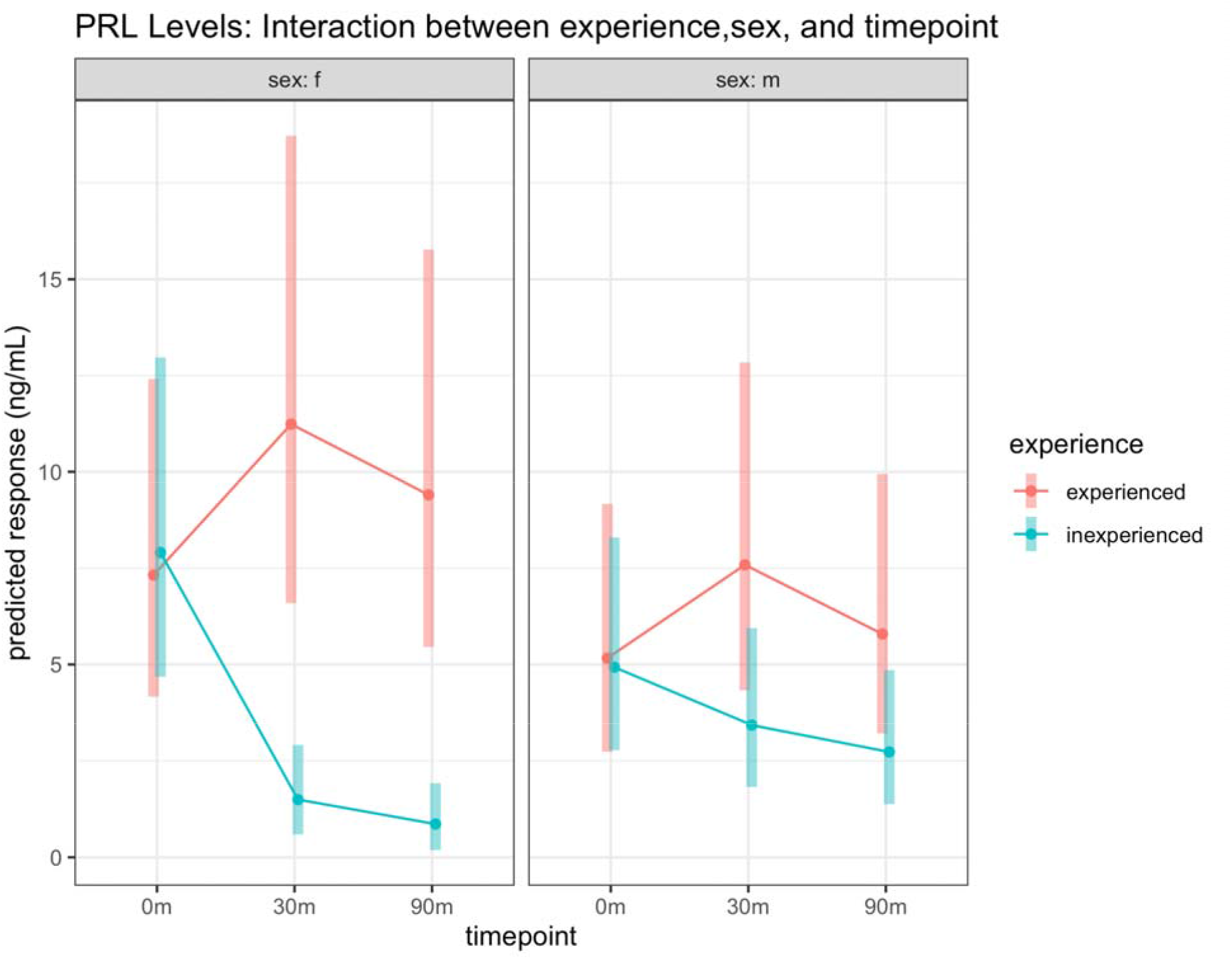
Interaction plot for estimated marginal mean PRL across parental experience, sex, and timepoint in non-actively nesting birds. Predicted responses from the mixed linear model, shown in concentration (ng/mL), for experienced (red) and inexperienced (blue) birds are across the stress series timepoints and sexes. The left plot shows trends in females and the right in males. Dots represent estimated marginal means and shaded areas represent 95% confidence intervals around these means. Plot produced using the emmeans package in R statistical language (Lenth, 2020).

